# Sulf2a controls Shh-dependent neural fate specification in the developing spinal cord

**DOI:** 10.1101/2020.08.20.258707

**Authors:** Cathy Danesin, Romain Darche-Gabinaud, Nathalie Escalas, Vanessa Bouguetoch, Philippe Cochard, Amir Al Oustah, David Ohayon, Bruno Glise, Cathy Soula

## Abstract

Sulf2a belongs to the Sulf family of extracellular sulfatases which selectively remove 6-O-sulfate groups from heparan sulfates, a critical regulation level for their role in modulating the activity of signalling molecules. Data presented here define Sulf2a as a novel player in the control of Sonic Hedgehog (Shh)-mediated cell type specification during spinal cord development. We show that Sulf2a depletion in zebrafish results in overproduction of V3 interneurons at the expense of motor neurons and also impedes generation of oligodendrocyte precursor cells (OPCs), three cell types that depend on Shh for their generation. We provide evidence that Sulf2a, expressed in a spatially restricted progenitor domain, acts by maintaining the correct patterning and specification of ventral progenitors. More specifically, Sulf2a prevents Olig2 progenitors to activate high-threshold Shh response and, thereby, to adopt a V3 interneuron fate, thus ensuring proper production of motor neurons and OPCs. We propose a model in which Sulf2a reduces Shh signalling levels in responding cells by decreasing their sensitivity to the morphogen factor. More generally, our work, revealing that, in contrast to its paralog Sulf1, Sulf2a regulates neural fate specification in Shh target cells, provides direct evidence of non-redundant functions of Sulfs in the developing spinal cord.

## INTRODUCTION

Sulf protein family, comprising Sulf1 and Sulf2, are heparan sulfate (HS) proteoglycan-editing enzymes that regulate a large number of signalling pathways^1,2^. These extracellular enzymes have been shown to present similar enzymatic activity, remodelling the 6-O-sulfation state of HS chains on the cell surface^3-5^. Thereby, Sulfs modulate HS binding to signalling molecules which leads to either promoting or inhibiting their signalling activities^1,2^. Sulf1 is the first discovered Sulf that was identified as a developmentally regulated protein, modulating Wnt signalling during somitogenesis in quail^6^. Since then, Sulf1 has been shown to control the signalling activity of other morphogen factors, including Sonic Hedgehog (Shh)^7-11^. In particular, Sulf1 behaves as a positive modulator of Shh signalling during spinal cord development, where it appears central in governing ventral neuron and oligodendrocyte specification^7-9^. How Sulf1 controls Shh signalling in the embryonic spinal cord has been partially elucidated^12^. In the ventral spinal cord, Shh signalling regulates generation of different types of neurons and glial cells which differentiate from distinct neural progenitors arrayed in domains along the dorso-ventral axis of the progenitor zone. Formation of these domains occurs early during neural tube patterning. Shh, emanating first from the notochord and later from the medial floor plate, spreads dorsally and induces expression of specific sets of transcription factors that trigger the formation of progenitor domains, each dedicated to generate a specific subtype of ventral neurons^13,14^. These domains emerge in a progressive manner and their order of appearance corresponds with their requirement for increasing concentrations and durations of Shh signalling^15,16^. This is illustrated by the sequential activation of Olig2 and Nkx2.2, two transcription factors expressed in the ventral-most progenitor domains. Nkx2.2, expressed in progenitors of the p3 domain and required for V3 interneuron specification, is located ventrally to progenitors of the pMN domain, which express Olig2 and generate somatic motor neurons (MNs)^13^. During patterning establishment, and in response to Shh, ventral neural progenitors first activate expression of Olig2. Afterwards, in response to a temporal increase in Shh signalling activity, the ventral-most progenitors activate Nkx2.2, a high-threshold Shh responsive gene. Nkx2.2, in turn, represses Olig2, thus leading to formation of the p3 and pMN non-overlapping domains. This is precisely at this step that Sulf1 is intervening. The enzyme, which is up-regulated in medial floor plate cells just prior to Nkx2.2 up-regulation, enhances Shh signalling activity by stimulating production of active forms of Shh from its source cells^7^. Noticeably, Shh stimulatory function of Sulf1 is reused later, at a time when pMN cells stop generating MNs and change their fate to generate oligodendrocyte precursor cells (OPCs). This temporal cell fate change depends on the formation of a novel source of Shh, named the lateral floor plate, that forms within the Nkx2.2-expressing progenitors of the p3 domain^12^. Again, Sulf1 must be up-regulated in these Shh-secreting cells so that these cells provide higher Shh signal to neighbouring pMN cells which, in turn, activate expression of Nkx2.2 that no longer represses Olig2 at this stage^17-20^. In this way, Sulf1 contributes to induce formation of a new progenitor domain, named the p* domain, populated by cells co-expressing Olig2 and Nkx2.2 fated to generate OPCs. Thus, in the developing spinal cord, the key role of Sulf1 is to change the inductive properties of Shh source cells to trigger high-threshold response to Shh.

Much less is known about the role of Sulf2 in regulating morphogen signalling. Sulf1 and Sulf2 have been proposed to display compensatory functions in many developmental processes^3,4,21-23^. However, whether Sulf1 and Sulf2 exert specific roles in the developing spinal cord remains to be addressed. In this tissue, in contrast to Sulf1 whose expression is restricted to medial and lateral floor plate cells, Sulf2 is expressed broadly in both floor plate cells and neural progenitors^7-11,24-26^. Comparison of Sulf1 and Sulf2 mouse knockout phenotypes led to the conclusion that the two enzymes act in a similar way at the level of Shh source cells to regulate Shh-dependent patterning at the time of OPC specification^11^. However, supporting the view that Sulf2 has a unique and additional function, conditional depletion of Sulf2 specifically in neural progenitors expressing Olig2 but not in Shh source cells is sufficient to impair gliogenesis in mouse^24^. Whether this function is linked to regulation of the Shh signal remains an open question.

In zebrafish, three Sulf members have been identified: Sulf1 and two Sulf2 paralogs, named Sulf2a and Sulf2b^27^. Here, we report that Sulf2a is the unique Sulf member whose expression is restricted to neural progenitors, then offering a simple context to study Sulf function specifically in Shh-responding cells of the developing spinal cord. We show that Sulf2a is required for proper generation of ventral neuronal and oligodendroglial cell subtypes and provide evidence that Sulf2a plays a key role in controlling neural progenitor identity. Overall, our data support a model whereby Sulf2a, by reducing the sensitivity of target cells to Shh, promotes low-threshold response to the morphogen over spinal cord development.

## RESULTS

### *Sulf2a* expression is restricted to neural progenitors of the developing spinal cord

As a first step to investigate the function of Sulf2 proteins in zebrafish, we analysed expression patterns of s*ulf2a* and s*ulf2b*. We first focused on two development stages: 24 hpf, while ventral neural pattern is established and neuronal production is ongoing and 36 hpf, corresponding to the time of p* domain formation and onset of OPC production in the ventral spinal cord. As previously reported^27^, we found that, at neurogenic stage (24 hpf), *sulf2b* is expressed in medial floor plate cells but also in adjacent neural progenitors of the p3 domain (Figure 1a, b). At the onset of gliogenesis (36 hpf), *sulf2b* signal, although weaker, was still detected in the medial floor plate and in adjacent cells which form to the lateral floor plate at that stage (Figure 1c, d). Thus, as previously reported for Sulf1^7^, *sulf2b* is expressed in Shh-producing cells of the developing spinal cord. Interestingly, we found a very different pattern of expression for *sulf2a* whose mRNA was detected in a broad domain of the ventral spinal cord at both neurogenic and gliogenic stages (Figure 1e-h). Noticeably, *sulf2a* mRNA was undetectable either in medial floor plate or in lateral floor plate cells but was found expressed in neural progenitors located at distance from Shh-producing cells (Figure 1f, h). Double detection of *sulf2a* and *shh* mRNAs further confirmed that *sulf2a* and *shh* domains of expression do not overlap neither at neurogenic (24 hpf) nor gliogenic (36 hpf) stages (Figure 1k, l). These data therefore revealed a unique property of *sulf2a* being exclusively expressed in neural progenitors.

**Figure 1:**
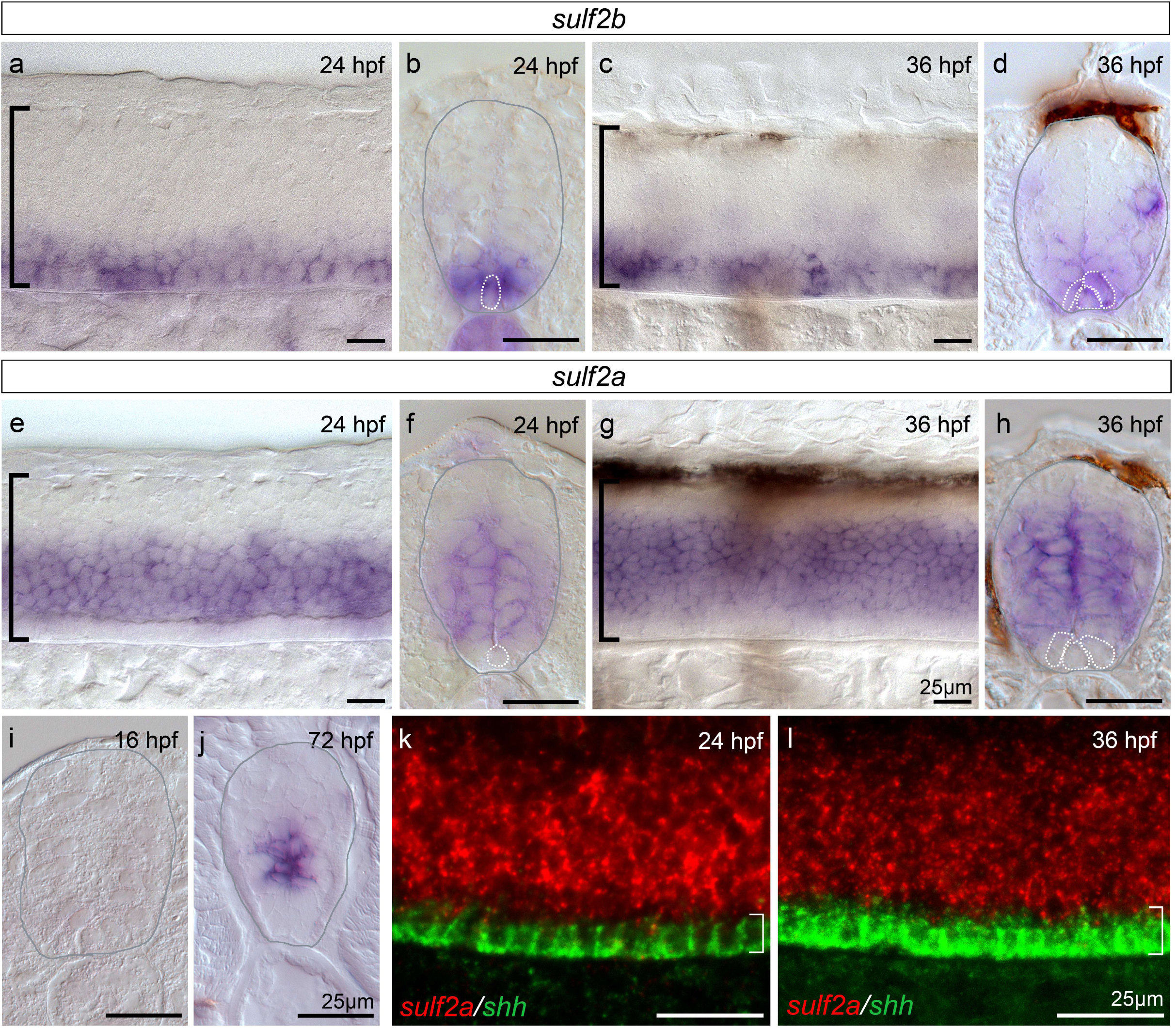
*Sulf2b* is expressed within Shh-producing cells whereas *Sulf2a* is restricted to neural progenitors. Here and in all subsequent panels, side views of embryos (a, c, e, g, k, l) are at the level of the trunk spinal cord, anterior to the left and dorsal to the top, and spinal cord transverse sections (b, d, f, h, i, j) are shown dorsal to the top. Developmental stages are indicated in each panel. **a-j**: Detection of *sulf2b* (a-d) and *sulf2a* (e-j) mRNAs by whole-mount *in situ* hybridisation. Images are representative examples of at least five embryos. Grey outlines (b, d, f, h, i, j) label the spinal cord edge. Brackets in a, c, e and g indicate the position of the spinal cord. Dotted lines in b, d, f and h delineate Shh-producing cells, i.e. the medial floor plate at 24 hpf and the medial floor plate + lateral floor plate at 36 hpf. Note that *sulf2a* mRNA is detected in neural progenitors but not in medial and lateral floor plate cells. **k-l**: Double detection of *sulf2a* (red) and *shh* (green) mRNAs by fluorescent *in situ* hybridisation, showing that *sulf2a* expression does not overlap the *shh*-expressing domain (white brackets).

We next investigated *sulf2a* expression at earlier and later stages of development. Ruling out the possibility that *sulf2a* plays a role in establishment of progenitor domains, we found that *sulf2a* is not yet expressed at 16 hpf (Figure 1i), while neural tube patterning has yet been established and neuronal production has already been initiated^7,28^. Much later, at the end of embryogenesis (72 hpf), s*ulf2a* expression was maintained in neural progenitors and was still excluded from Shh-producing cells (Figure 1j), thus indicating that *sulf2a* remains expressed in neural progenitors after the onset of neuronal production and all along gliogenesis.

Together, these results, showing that *sulf2a* is the only *sulf* gene activated specifically in neural progenitors open the possibility that this Sulf member plays a specific role in regulating signal reception at the surface of spinal progenitors as they generate neurons and glial cells.

### Sulf2a controls generation of both V3 interneurons and OPCs

Our data showing that *sulf2a* is activated after initiation of neuronal production and persists along gliogenesis, prompted us to examine its function in regulating generation of V3 interneurons and OPCs produced by ventral neural progenitors^29,30^. To address the role of Sulf2a, we first used a morpholino (MO) knockdown approach, using a translation (*sulf2a*MO^ATG^) and a splice (*sulf2a*MO^splice^) blocking morpholino. As similar results were obtained using either MO in all experiments, they are referred to as *sulf2a*MO in the text. We first assessed V3 interneuron generation by detecting *sim1a* expression as a specific marker of these neurons originating from the p3 domain^29,31^. At 48 hpf, as neuronal production has been completed, we found that the number of *sim1a*+ cells was significantly higher in *sulf2a*MO-injected embryos compared to the control MO (ctrlMO)-injected ones, with more than a 20% increase in the number of V3 interneurons in *sulf2a* morphants (Figure 2a-c, f). To confirm these observations, we generated a *sulf2a* mutant zebrafish line using the CRISPR/Cas9 system. We targeted the hydrophilic domain (HD) region of *Sulf2a*, known to be required for enzymatic activity and substrate recognition^3,32-34^. We obtained an indel mutation in the coding region which leads to a premature translation termination and a predicted protein missing functional HD domain. Zebrafish carrying homozygous *sulf2a* mutation (*sulf2a*-/-) showed no overt phenotype, are viable, fertile, and produce offsprings that show no obvious morphological defects. Confirming the requirement of Sulf2a to restrict V3 interneuron production, we found that the number of *sim1a*+ neurons increased by 25% in *sulf2a*-/- embryos compared to wild-type siblings (Figure 2d-f). We conclude from these data that Sulf2a plays a role in limiting production of V3 interneurons.

**Figure 2:**
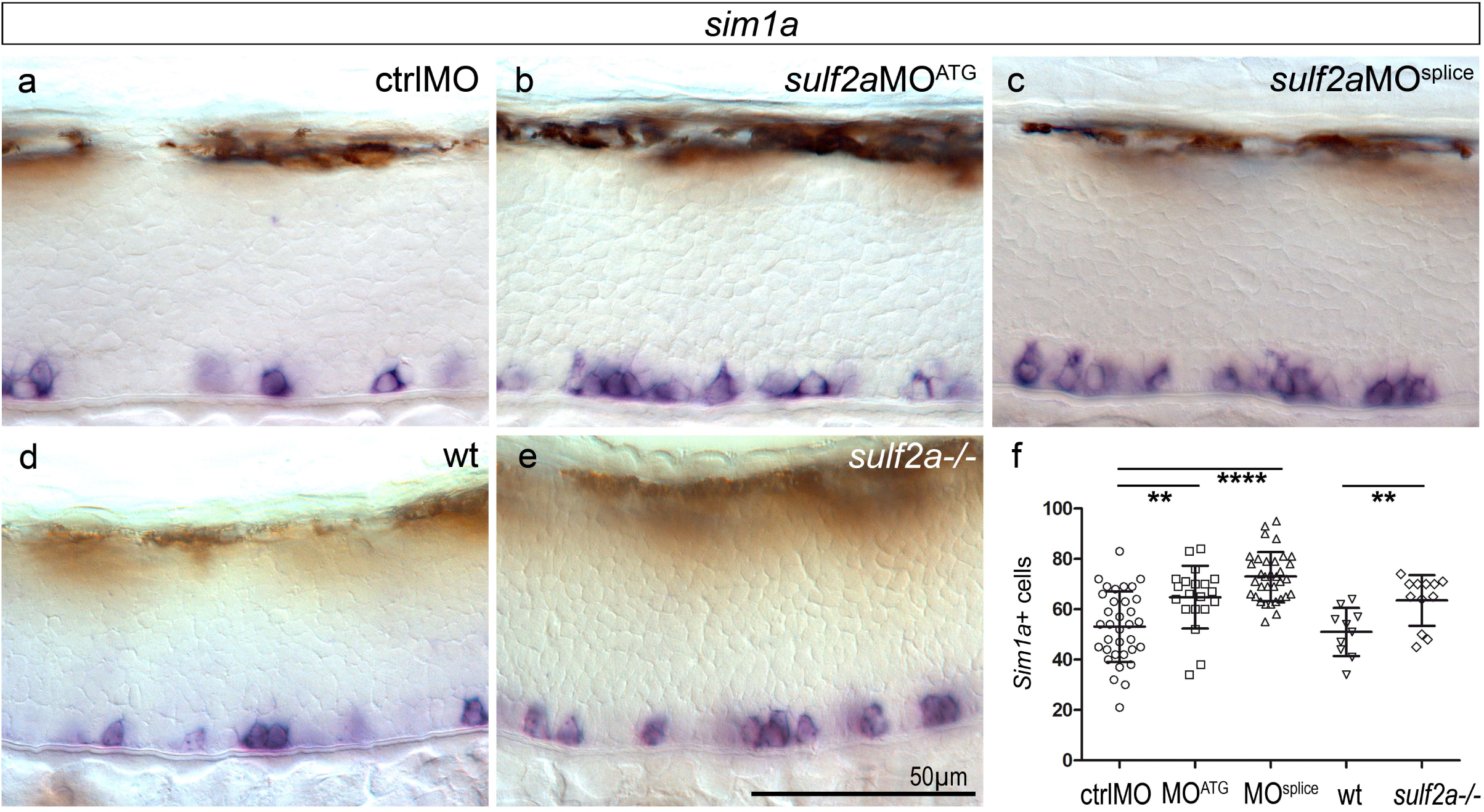
V3 interneuron production is increased upon *Sulf2a* depletion. Side views of 48 hpf embryos. **a-f**: Detection (a-e) and quantification (f) of *sim1a*-expressing neurons in embryos injected with ctrlMO (a, n=34), *sulf2*aMO^ATG^ (b, n=20) or *sulf2a*MO^splice^ (c, n=34) and in wild-type (d, n=10) or *sulf2a*-/- (e, n=12) embryos from two independent experiments in each case. Datasets were compared with Mann Whitney’s test (two-tailed). Data are presented as mean number of cells per embryo ± s.d (**p<0.01, ****p<0.0001).

*Sulf2*a expression being maintained at 36 hpf, the time of OPC specification^30^, we next asked whether Sulf2a might also play a role in the control of OPC generation. To identify OPCs, we used either Tg(*olig2:*GFP) or Tg(*olig2*:DsRed2) transgenic lines in which fluorescently-labelled OPCs can be identified by their morphology and position^35,36^. Due to expression of *olig2* in progenitors of the pMN domain as well as perdurance of GFP or DsRed2 in newly generated MNs^35-37^, these reporter lines however do not allow distinguishing OPCs from other cell types in the ventral spinal cord. We bypassed this problem by performing immunodetection of Sox10, the earliest specific marker of oligodendroglial cells^37^. At 48 hpf, when OPC generation has just began, we found *olig2:*GFP+ or *olig2:*DsRed2+ cells co-expressing Sox10 in the ventral spinal cord of both ctrlMO-injected and wild-type embryos (Figure 3a, d). At identical stage, the number of OPCs was significantly reduced either in *sulf2a*MO-injected or in *sulf2a*-/- embryos, with a decrease of more than 45% and 33% compared to controls (Figure 3b, c, e, f). Thus, Sulf2a activity is also required for proper generation of OPCs but, instead of limiting the number of cells as observed for V3 interneurons, the enzyme contributes to promote OPC production.

**Figure 3:**
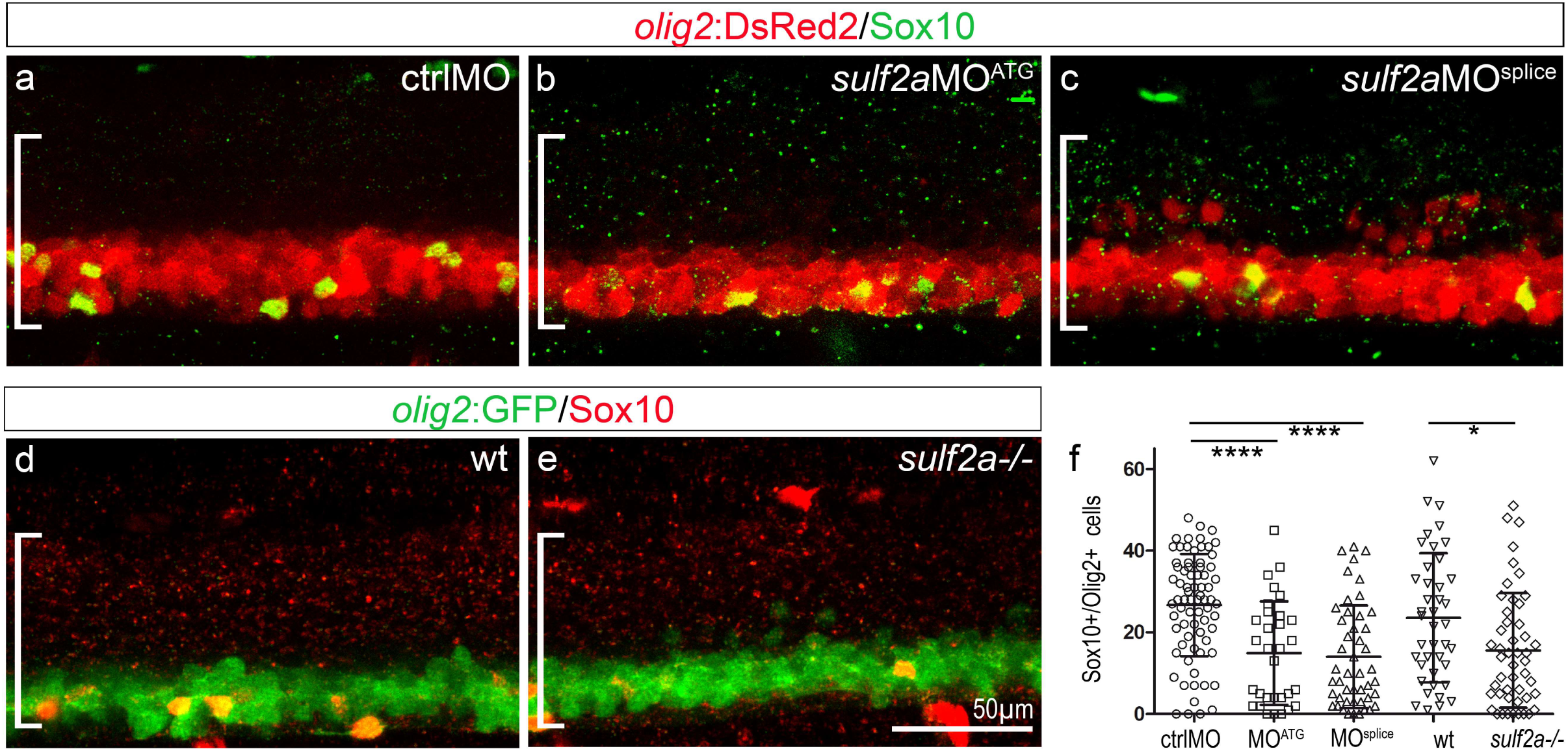
Sulf2a depletion impairs OPC development. Side views of 48 hpf embryos. **a-f**: Detection (a-e) and quantification (f) of OPCs by immunodetection of Sox10 (green in a-c, red in d, e) in Tg(*olig2*:DsRed) embryos (red in a-c) injected with ctrlMO (a, n=73), *sulf2*aMO^ATG^ (b, n=31) or *sulf2a*MO^splice^ (c, n=47) and wild-type (d, n=10) or *sulf2a*-/- (e, n=48) Tg(*olig2*:GFP) embryos (green in d and e) from 9 and 6 independent experiments, respectively. Datasets were compared with Mann Whitney’s test (two-tailed). Data are presented as mean number of cells per embryo ± s.d (*p< 0.05, ****p<0.0001).

Together, these results, showing that Sulf2a limits production of V3 interneurons during neurogenesis but promotes generation of OPCs at initiation of gliogenesis, reveal specific and differential functions of Sulf2a in controlling generation of ventral neural cell subtypes.

### Sulf2a controls the identity of ventral neural progenitors

We next focused on the question of how the same enzyme can fulfil differential regulatory functions to ensure production of a correct number of neural derivatives. *Sulf2a* being expressed in neural progenitors, we postulated that the enzyme might be involved in regulating progenitor patterning and subsequent cell diversification. We started analysing progenitor domain organisation at 24 hpf while neurons are being produced. We focused on the p3 and pMN domains characterised by the expression of Nkx2.2a and Olig2, respectively. Nkx2.2a expression was assessed detecting mEGFP in the Tg(*nkx2.2a*:mEGFP) line^36^, while Olig2 expression was monitored by detecting the *olig2* mRNA using fluorescent RNA *in situ* hybridisation. Immunodetection of Sox2, a generic marker of neural progenitors^38,39^, was also performed to distinguish these cells from newly generated neurons. Analysis of ctrlMO-injected embryos revealed *nkx2.2a*:mEGFP+ progenitors organised in a row of one to two cells in the ventral progenitor zone and comparison with *olig2* mRNA staining showed that *olig2* expression defined the dorsally adjacent non-overlapping progenitor domain (Figure 4a-a”), thus reflecting ventral patterning establishment. Similar analysis performed in *sulf2a*MO-injected embryos showed that the *nkx2.2a*:mEGFP+ domain extended dorsally, covering rows of two to three cells, reflecting a more than 35% broadening of the p3 domain compared to controls (Figure 4b, b”, c, c”, d). Thus, in agreement with the excess of V3 interneurons observed in Sulf2a-depleted embryos, p3 progenitor number is increased in these embryos. By contrast, *olig2* expression was found markedly reduced in *sulf2a*MO-injected embryos (Figure 4b’-c’’). Counting of *olig2*+ progenitors indicated a more than 59% decrease in *sulf2a* morphants compared to control embryos (Figure 4e). Accordingly, the number of MNs, assessed by immunostaining of Islet1/2 in Tg(*olig2*:DsRed2) embryos, was reduced by 20-30% in 48hpf *sulf2a-*depleted embryos (Supplementary Figure 1). Together, these data indicated that Sulf2a is part of the regulatory process that controls dorsal boundary positioning of the Nkx2.2a/p3 domain and thus contributes to maintain the Olig2/pMN domain over the period of neuronal production.

**Figure 4:**
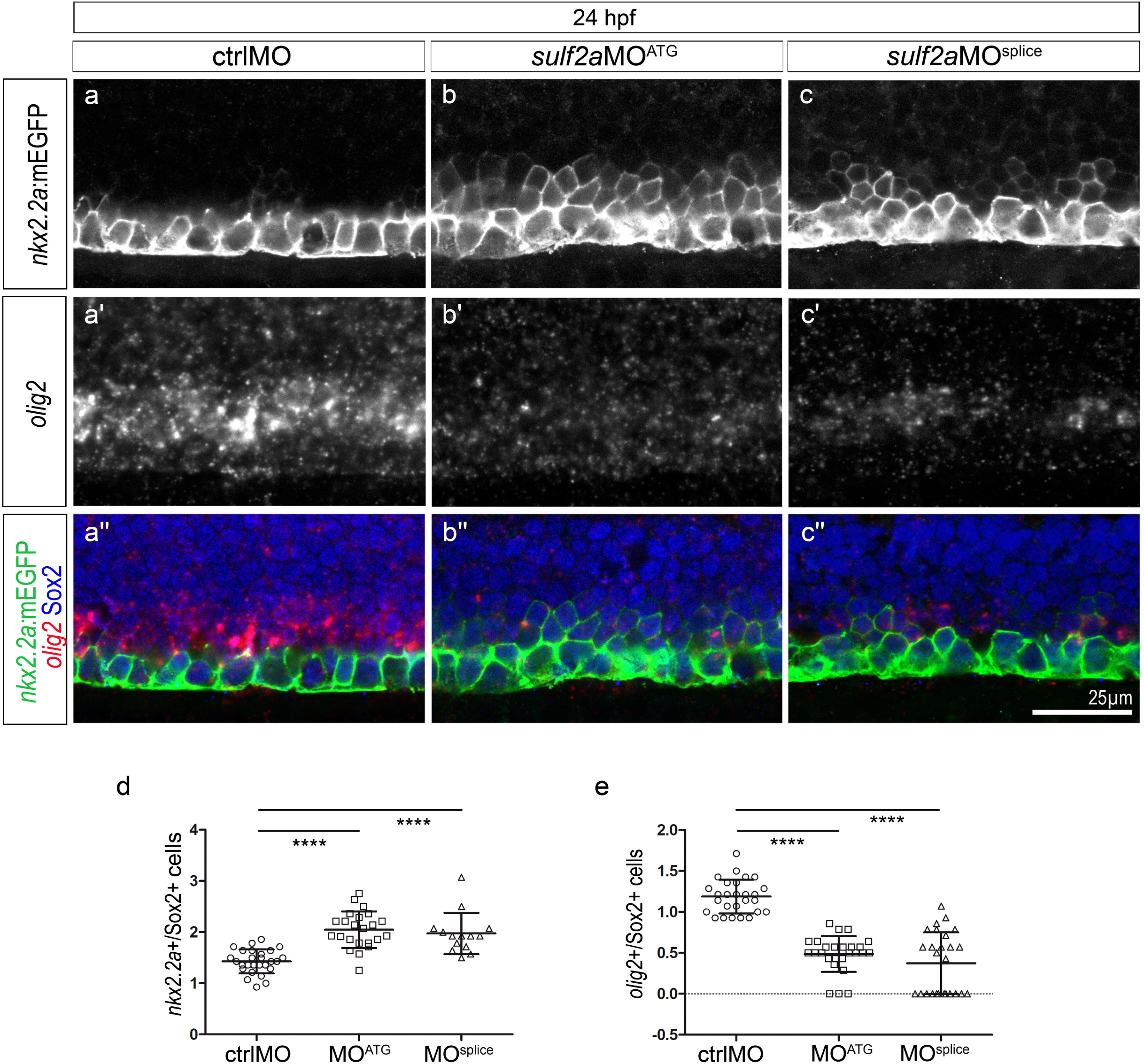
The Nkx2.2a/p3 domain is dorsally expanded at the expense of the Olig2/pMN domain in 24 hpf *sulf2a*-depleted embryos. Side views of 24 hpf embryos. **a-c’’**: Double detection of *olig2* mRNA and Sox2 in Tg(*nkx2.2a*:mEGFP) embryos injected with ctrlMO (a-a”), *sulf2a*MO^ATG^ (b-b”) or *sulf2a*MO^splice^ (c-c”). Vertical sets present successively the *nkx2.2a*:mEGFP signal (green in a’’-c”), the *olig2* mRNA staining (red in a’’-c’’) and the merged image together with Sox2 staining (blue in a”-c”). Note the dorsal expansion of the *nkx2.2a:*mEGFP signal associated with a strong decrease in the *olig2* signal in *sulf2a* morphant embryos. **d, e**: Quantification of *nkx2.2a:*mEGFP+/Sox2+ (d) and *olig2*+/Sox2+ (e) progenitors in embryos injected with ctrlMO (n=27), *sulf2a*MO^ATG^ (n= 14) and *sulf2a*MO^splice^ (n= 23) from two independent experiments. Datasets were compared with Mann Whitney’s test (two-tailed). Data are presented as a mean number of cells along the dorso-ventral axis per embryo ± s.d (****p<0.0001).

We next examined neural patterning at the onset of gliogenesis (36 hpf). As expected, we found that, in control embryos, the *nkx2.2a*:mEGFP+ progenitor domain was broader than that observed at the neurogenic period (compare Figures 4 and 5) and partially overlapped the *olig2-*expressing domain (Figure 5a”, a”’), reflecting formation of the p* domain at this stage. As observed at neurogenic period, *sulf2a* depletion caused dorsal expansion of the *nkx2.2a*:mEGFP+ domain, with a more than 25% increase in the number of *nkx2.2a*:mEGFP+ progenitors compared to controls (Figure 5b, b’’, b’’’, c”, c’”, d), thus indicating that the dorsal boundary position of the Nkx2.2a-expressing domain is also affected by Sulf2a depletion at the time of patterning rearrangement. Counting of *olig2*+ progenitors in *sulf2a* morphants showed that their number decreased by 40-50%, a reduction still significant but less pronounced than that observed at neurogenic stage (Figure 5b’, c’, e). Noticeably and in agreement with the loss of repressive activity of Nkx2.2a on *olig2* expression at this stage, no significant difference was found in the number of progenitors co-expressing *nkx2.2a*:mEGFP and *olig2* between control and *sulf2a* morphant embryos (Figure 5a”-c”’, f). The logical corollary of this latter observation is that, upon *sulf2a* depletion, the observed reduction in the number of Olig2 progenitors should be due to a specific decrease in the number of progenitors expressing Olig2 but not Nkx2.2a. Accordingly, quantification of these cells showed a decrease by more than 60% in *sulf2a-*depleted embryos (Figure 5 a”-c”’g). Thus, at gliogenic stage, Sulf2a depletion causes dorsal expansion of the Nkx2.2a-expressing domain, does not affect generation of the p* domain but leads to a drastic loss of the dorsally located progenitors that express Olig2 but not Nkx2.2a. From these data, we conclude that, at stage of patterning rearrangement, Sulf2a function is again required to position dorsal boundary expression of the Nkx2.2a domain, but also to ensure that a population of Olig2-expressing progenitors that do not activate Nkx2.2a are generated.

**Figure 5:**
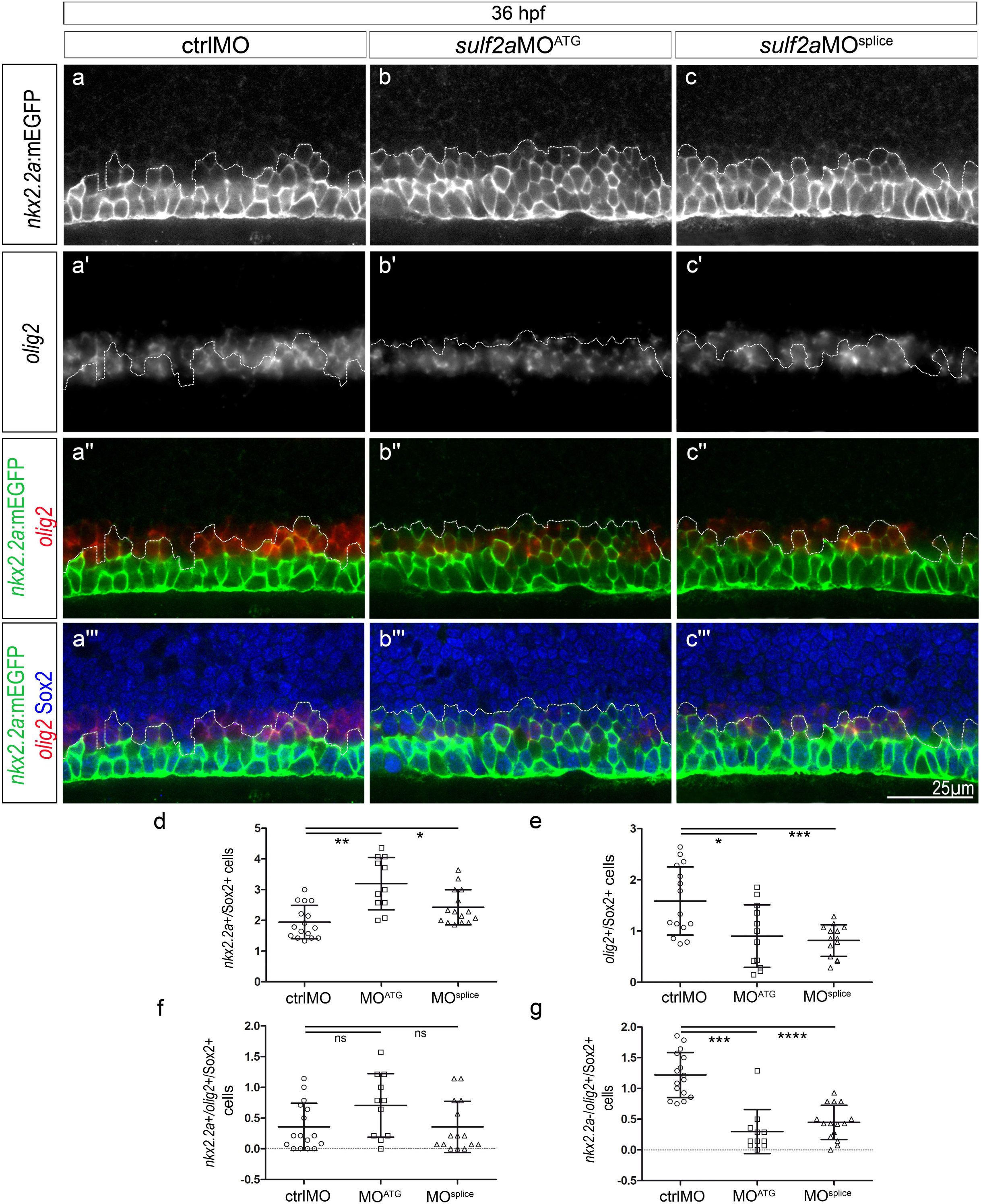
*Sulf2a* depletion does not affect formation of the Nkx2.2a/Olig2-co-expressing p* domain but reduces the number of progenitors that only express Olig2 at the onset of gliogenesis. Side views of 36 hpf embryos. **a-c’’’**: Double detection of *olig2* mRNA and Sox2 in Tg(*nkx2.2a*:mEGFP) embryos injected with ctrlMO (a-a”’), *sulf2a*MO^ATG^ (b-b”’) or *sulf2a*MO^splice^ (c- c”’). Vertical sets present successively the *nkx2.2a*:mEGFP signal (green in a’’-c”, a”’-c”’), the *olig2* mRNA staining (red in a”-c”, a”’-c”’), the merged image of *nkx2.2a*:mEGFP and *olig2* signals and the merged image together with Sox2 staining (blue in a”’-c”’). Dotted line represents the dorsal limit of *nkx2.2a*:mEGFP-expressing domain. **d-g**: Cell quantification performed in embryos injected with ctrlMO (n=16), s*ulf2a*MO^ATG^ (n= 11) and *sulf2a*MO^splice^ (n=15) from three independent experiments. Cell counts in d and e correspond to the population of Sox2+ progenitors expressing either *nkx2.2a*:mEGFP (d) or *olig2* (e). Quantifications in f and g correspond to Sox2+ progenitors that either co-express *olig2* and *nkx2.2a*:mEGFP (f) or express *olig2* mRNA but not *nkx2.2a*:mEGFP (g). Except in e where datasets were compared with Student’s *t*-test (unpaired two-tailed), datasets were compared with Mann Whitney’s test (two-tailed). Results are presented as a mean number of cells along the dorso-ventral axis per embryo ± s.d (*p<0,05, ** p<0,005, ***p<0.001,****p<0.0001).

Together, these data highlight the key role of Sulf2a in assigning and maintaining neural progenitor identity at neurogenic and gliogenic stages.

### Sulf2a regulates allocation of oligodendrocyte lineage cell fates

In zebrafish, newly specified OPCs, all expressing Olig2, represent a heterogeneous cell population in regard to Nkx2.2a expression and fate. While some OPCs express Nkx2.2a and differentiate rapidly as oligodendrocytes, others, that do not express Nkx2.2a, remain as non-myelinating OPCs ^40,41^. Hereafter we will refer to these two OPC subsets as myelinating and non-myelinating OPCs, respectively. Whether these two OPC subsets emerge from Olig2 neural progenitors already distinguishable by differential Nkx2.2a expression remained an open question. Our data, highlighting the requirement of Sulf2a specifically for maintaining progenitors expressing only Olig2, led us to ask whether Sulf2a contributes specifically to generate the non-myelinating OPC subset. We thus examined the consequences of Sulf2a depletion on the development of the two distinct OPC populations by performing immunodetection of the pan-OPC marker Sox10 in Tg(*nkx2.2a*:mEGFP; *olig2:*DsRed2) embryos in which the two OPC subtypes can be distinguished by differential reporter expression^36^. As expected, at early gliogenic period (48 hpf), we found that the global population of OPCs, i.e. the Sox10+/*olig2:*DsRed2+ cells, was reduced in number either in *sulf2a*MO-injected or in *sulf2a*-/- embryos, decreasing by 40% compared to controls (Figure 6a, b, c, d, e). We next considered separately the myelinating (*olig2*:DsRed2+/*nkx2.2a*:mEGFP+) and the non-myelinating (*olig2*:DsRed2+/*nkx2.2a*:mEGFP-) OPC subtypes. Quantification of these cells pointed to a decrease in number of both OPC populations but to a different extent. In *sulf2a* morphants, non-myelinating OPCs were found decreased in number by 64% (Figure 6a-b”, g) while the myelinating population was reduced only by 31% compared to controls (Figure 6a-b”, f). A similar tendency was obtained analysing the two OPC subsets in *sulf2a*-/- embryos (Figure 6c-d”, f, g). Thus, the impaired development of OPCs observed in Sulf2a-depleted embryos is mostly attributable to a decreased number of OPCs that do not express Nkx2.2a.

**Figure 6:**
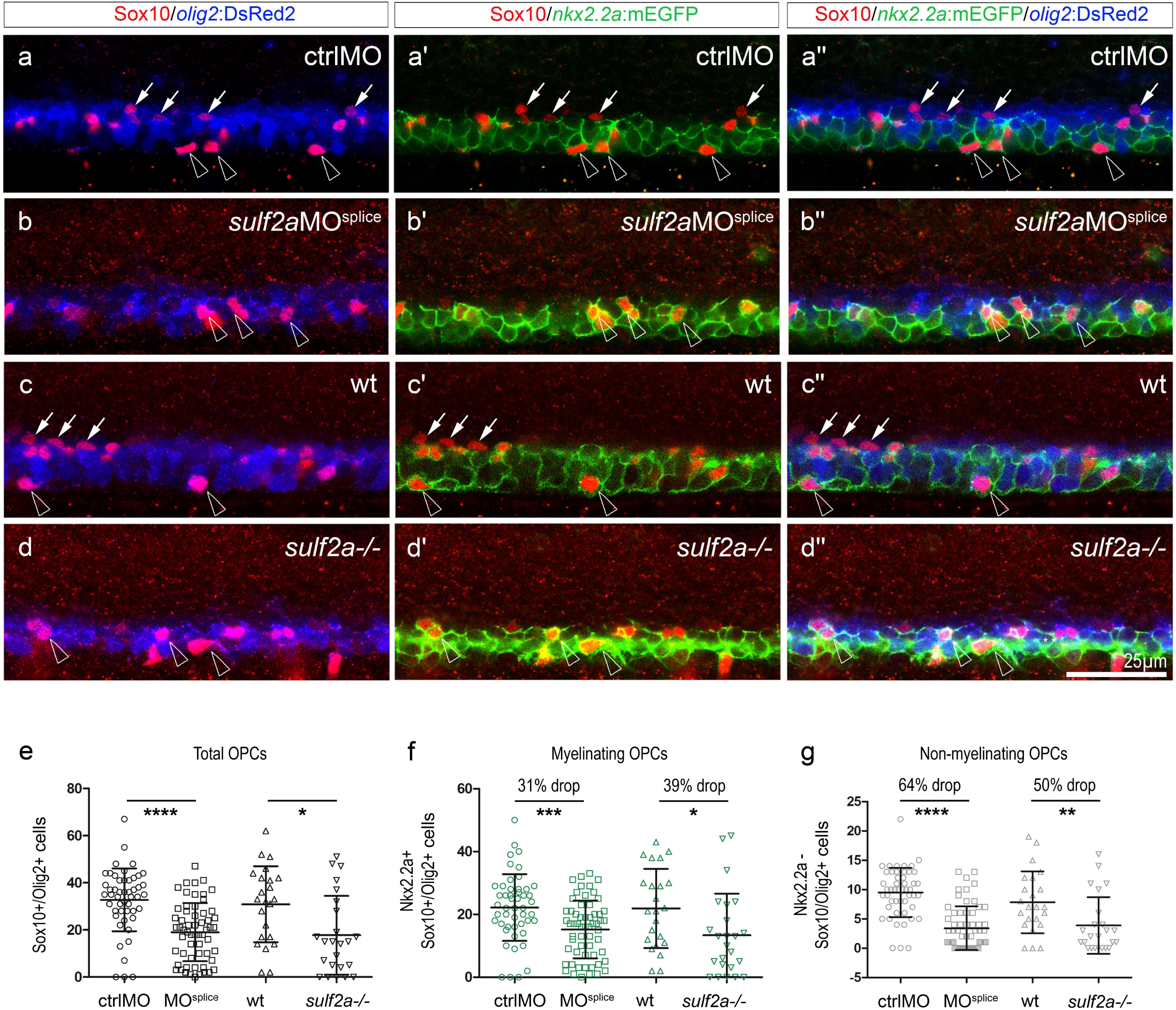
*Sulf2a* depletion causes a preferential deficit of non-myelinating OPCs. Side views of 48 hpf embryos. **a-d’’**: Immunodetection of Sox10 in Tg(*nkx2.2a*:mEGFP; *olig2*:DsRed2) embryos injected with ctrlMO (a-a”) or *sulf2a*MO^splice^ (b-b”) and wild-type (c-c”) or mutant for *sulf2a* (d-d”). Horizontal sets present successively the Sox10 staining (red) in combination with the *olig2*:DsRed2 signal (blue, a-d), with the *nkx2.2a*:mEGFP signal (green, a’-d’) and the merged image of the three signals (a”-d”). White arrows point to non-myelinating OPCs (*Nkx2.2a*:mEGFP-/*olig2*:DsRed2+/Sox10+ cells) while open arrowheads indicate myelinating OPCs (*Nkx2.2a*:mEGFP+/*olig2*:DsRed2+/Sox10+ cells). **e-g**: Quantification of OPCs in embryos injected with ctrlMO (n=70) or *sulf2a*MO^splice^ (n=59) and in wild-type (n=23) or *sulf2a*-/- (n=24) embryos from five and three independent experiments, respectively. Cell counts in e correspond to the total number of OPCs co-expressing Sox10 and *olig2*:DsRed2. Quantifications of *nkx2.2a*:mEGFP+/myelinating OPCs (f) and *nkx2.2a*:mEGFP-/non-myelinating OPCs (g) were presented separately. Datasets were compared with Mann Whitney’s test (two-tailed). Data are presented as mean number of cells per embryo ± s.d (*p<0.05, **p<0.01, *** p<0.0005, ****p<0.0001).

These results, showing that Sulf2a plays a prominent role in promoting the development of non-myelinating OPCs, support the view that Sulf2a contributes to the early segregation of the two OPC lineages by preserving a pool of progenitors that only express Olig2.

### Sulf2a expression in neural progenitors prevents them to activate high-threshold response to Shh

Our results, pointing to a regulatory function of Sulf2a on limiting the dorsal extent of Nkx2.2a expression, highlight the key role of Sulf2a in maintaining Olig2-expressing progenitors at neurogenic and gliogenic stages. Regulation of *olig2* and *nkx2.2a* expression is known to depend on different levels of Shh signalling in zebrafish, *nkx2.2a* requiring higher doses to be induced ^7,37,42,43^. Thus, an attractive hypothesis was that Sulf2a acts by preventing activation of high Shh signalling in ventral neural progenitors. According to this possibility, *nkx2.2a*+ cells should not express *sulf2a*. To test this, we compared *sulf2a* and *nkx2.2a* expression domains at 24 and 36 hpf performing *sulf2a* fluorescent *in situ* hybridisation in Tg(*nkx2.2a*:mEGFP) embryos. We found that, at each stage, *sulf2a* and *nkx2.2a*:mEGFP expression defined two distinct domains in the ventral developing spinal cord (Figure 7a-d), thus confirming that expression of *sulf2a* is excluded from *nkx2.2a*-expressing cells. Noticeably, ventral boundary of the *sulf2a*+ domain was found to abut dorsal boundary of the *nkx2.2a*:mEGFP*+* domain at each stage (Figure 7a-d). These observations suggested that, at 24 hpf, *sulf2a* expression overlaps the entire Olig2+ domain while it is not the case at 36 hpf, since, at that stage, the dorsal-most Nkx2.2a+ cells are known to co-express Olig2. To clarify this, we analysed expression of *sulf2a* mRNA at 36 hpf in Tg(*olig2*:GFP) embryos and found expression of *sulf2a* in a subset of Olig2 progenitors (Figure 7e, f). *Sulf2a* expression being excluded from the Nkx2.2a-expressing domain, we conclude that these cells correspond to Olig2 progenitors that are not included in the p* domain. Overall, these data, showing that *sulf2a* is excluded from progenitors that respond to high Shh signalling activity (Nkx2.2a+ cells) but expressed in progenitors that do not activate high-threshold response to Shh (Olig2+ cells), support the view that Sulf2a might act at the surface of Olig2+ progenitors to reduce their sensitivity to Shh signalling. They moreover highlight differential expression of *sulf2a* in Olig2 progenitors at gliogenic stage.

**Figure 7:**
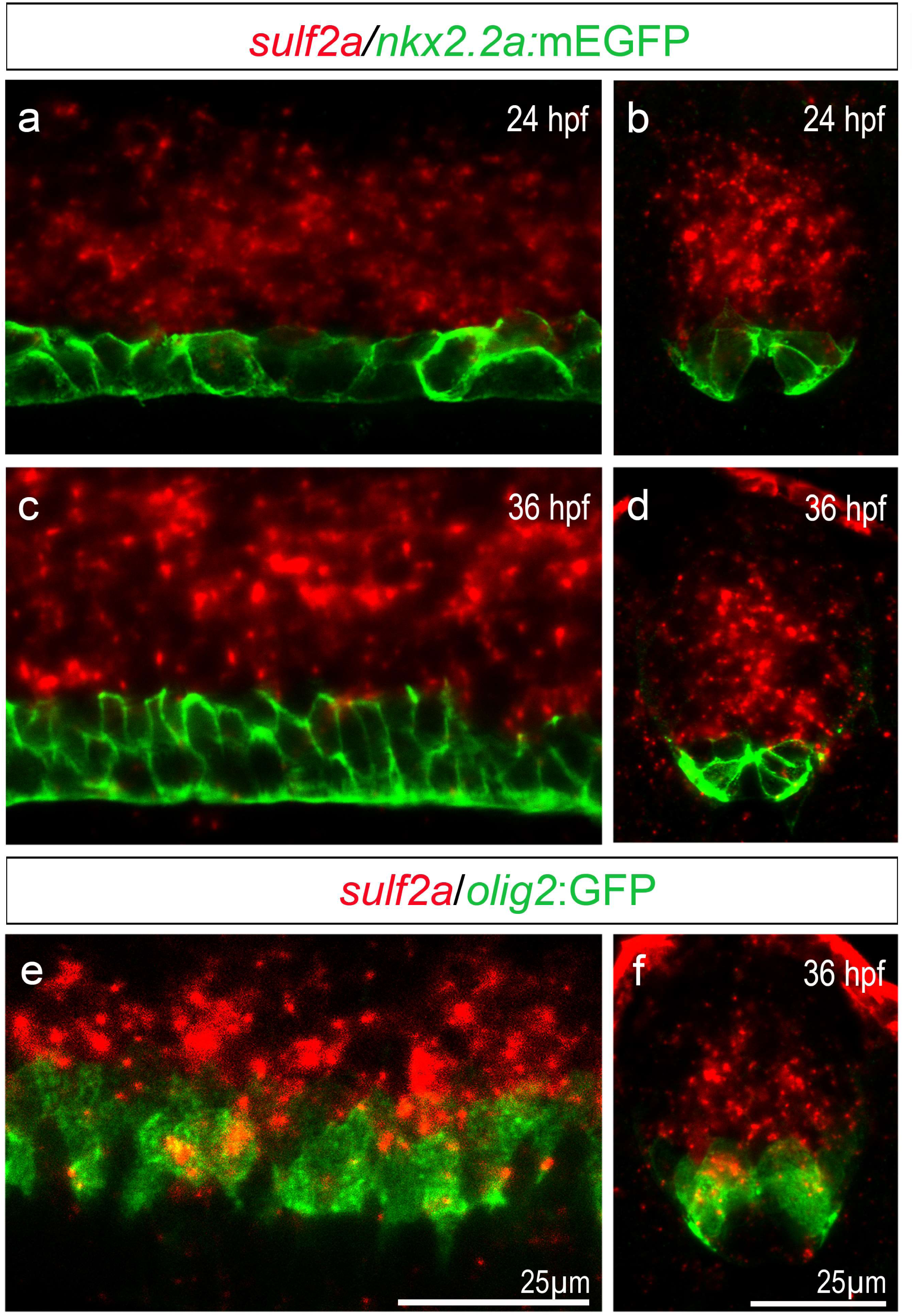
*Sulf2a* is excluded from Nkx2.2a-expressing progenitors but expressed, at gliogenic stage, in a subset of Olig2 progenitors. Side views (a, c, e) and transverse sections (b, d, f) of 24 hpf (a, b) and 36 hpf (c-f) embryos. a-f: Detection of *sulf2a* mRNA (red) in Tg(*nkx2.2a*:mEGFP) embryos (green, a-d) or in Tg(*olig2:*GFP) embryos (green, e, f). Images show representative examples of at least three embryos. Note that *sulf2a* expression dorsally abuts the *nkx2.2a*:mEGFP-expressing domain both at 24 and 36 hpf (a-d) and is heterogeneously expressed within the population of Olig2 progenitors at 36 hpf (e, f).

An attractive hypothesis was thus that Sulf2 restricts the range of high-threshold response to Shh along the ventral to dorsal axis of the progenitor zone. To test this possibility, we used Tg(*GBS-ptch2*:EGFP) reporter line in which cells responding to Shh can be visualized by EGFP expression^44^. At 24 hpf, in control embryos, *GBS-ptch2*:EGFP expression was detected in the ventral neural tube in medial floor plate cells as well as adjacent progenitors that both express Sox2 (Figure 8a, b). We interpreted the EGFP signal in medial floor plate cells as perdurance of the reporter protein since, while responsible for medial floor plate induction at earlier stages, Shh signalling is known to be inactive in these cells at 24 hpf^45,46^. Moreover, EGFP was not detected at distance from the Nkx2.2a/p3 domain, indicating that the EGFP signal only allowed detecting the highest levels of Shh signalling. Having this in mind, we next examined EGFP expression in *sulf2a*MO-injected embryos. We found that the EGFP expression domain was enlarged and cell quantification confirmed a slight but significant increase in the number of EGFP+/Sox2+ cells compared to controls (Figure 8c-e), thus validating the role of Sulf2a in limiting the dorsal extent of high-threshold Shh signalling activity. Importantly, we found that *shh* expression was not modified in Sulf2a-depleted embryos (Figure 8 f-m), indicating that the influence of Sulf2a on Shh signalling activity does not depend on transcriptional regulation of the morphogen factor. These data, showing that Sulf2a activity is required to limit the dorsal extent of high-threshold Shh response, highlight a functional relationship between Sulf2a and Shh and bring support to the view that the enzyme acts by reducing the sensitivity of target cells to the Shh signal.

**Figure 8:**
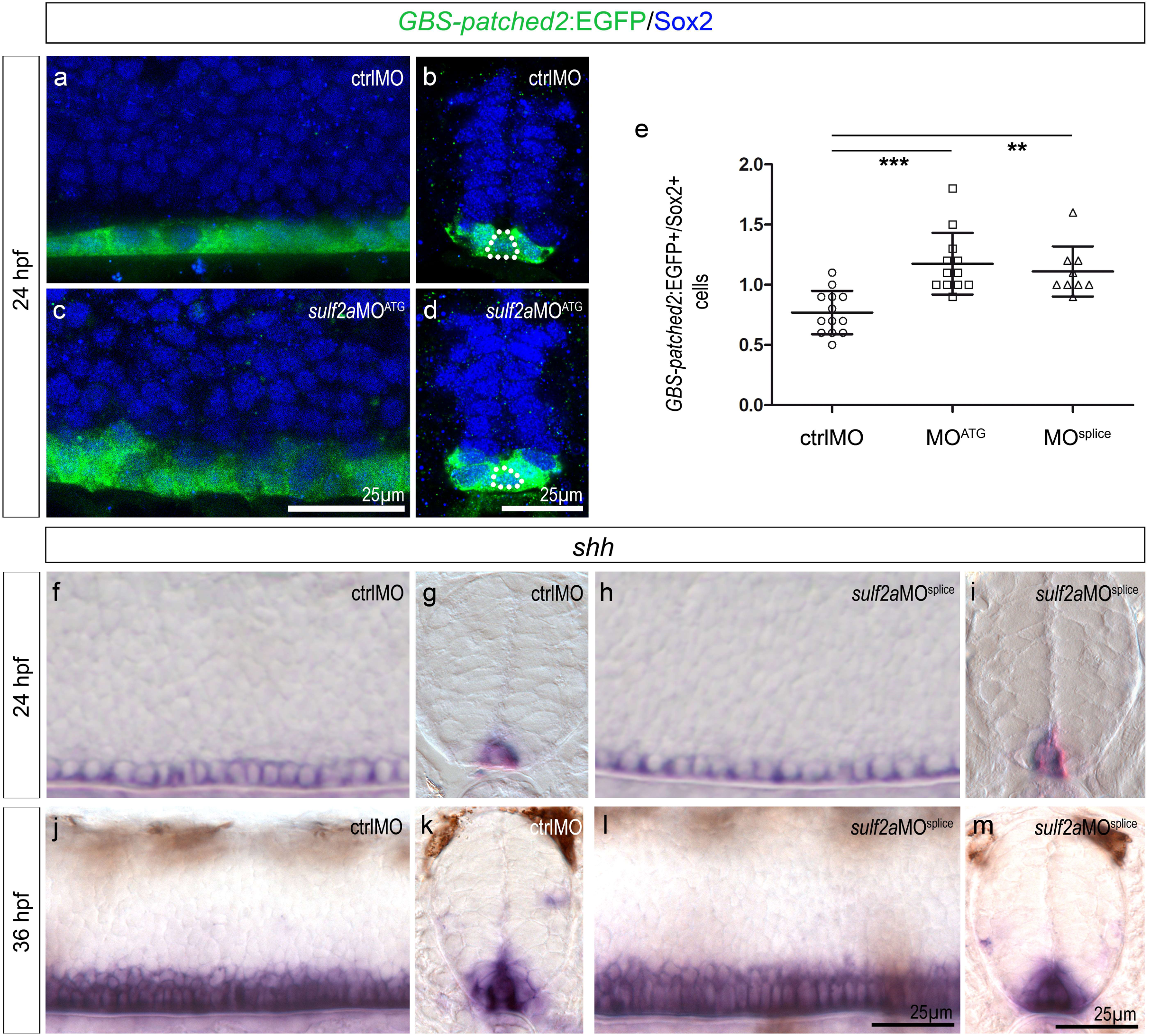
*Sulf2a* restricts dorsal extent of the high-threshold Shh response without affecting expression of *shh*. Side views (a, c, f, h, j, l) and transverse sections (b, d, g, i, k, m) of 24 hpf (a-i) and 36 hpf (j-m) embryos. a-d: Immunostaining of Sox2 in Tg(*GBS-ptch2*:EGFP) embryos injected with ctrlMO (a, b) or *sulf2a*MO^ATG^ (c, d). Dotted lines in b and d delineate the medial floor plate. Note the dorsal expansion of the *GBS-ptch2*:EGFP signal in *sulf2a* morphants. e: Quantification of *GBS-ptch2*:EGFP+/Sox2+ cells in embryos injected with ctrlMO (n=13), *sulf2a*MO^ATG^ (n=12) or *sulf2a*MO^splice^ (n=9) from two independent experiments. Datasets were compared with Mann Whitney’s test (two-tailed). Data are presented as mean number of cells along the dorso-ventral axis on each side of the lumen per embryo ± s.d (***p< 0.0005, ** p<0.005). f-m: Detection of *shh* mRNA in embryos injected with ctrlMO (f, g, j, k) or *Sulf2a*MO^splice^ (h, i, l, m).

## DISCUSSION

In the ventral spinal cord, generation of distinct neural cell types is a spatio-temporally regulated process that mainly depends on the morphogenetic activity of Shh. During development, neural progenitors are exposed to varying concentrations of Shh which drive them to adopt specific transcriptional identities and eventually generate a particular neuronal or glial cell fate. We show here that Sulf2a contributes to assign specific neural identities to Shh-responding progenitors. We provide evidence that expression of *sulf2a* in Olig2 progenitors is essential to prevent highest levels of Shh signalling within these cells, thus allowing them to orient toward the production of, first MNs and, later on, of a particular subtype of OPC. Our data support a model in which Sulf2a spatially restricts expression of high-threshold Shh responsive genes to cells closest to the source of Shh (Figure 9).

**Figure 9:**
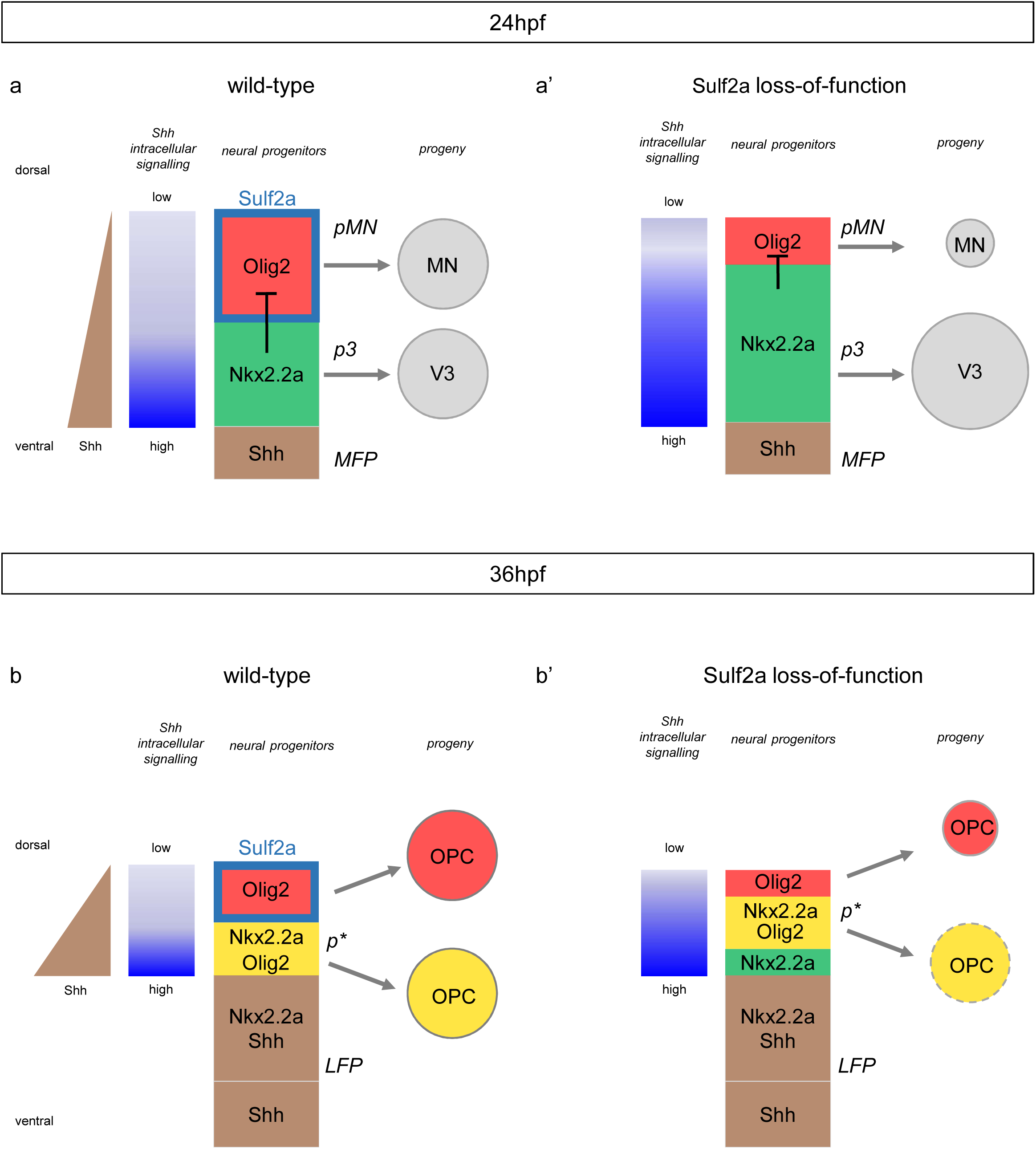
Model for Sulf2a function. Scheme showing Sulf2a and Shh-dependent gene expression in ventral neural progenitors during neuronal production (24 hpf, a, a’) and at the onset of gliogenesis (36 hpf, b, b’) in wild-type (a, b) and *sulf2a* depleted (a’, b’) contexts. In 24 hpf wild-type embryos (a), the ventral-most progenitors of the p3 domain (green), adjacent to the medial floor plate (MFP, brown), express the high-threshold Shh responsive gene Nkx2.2a and generate V3 interneurons. Sulf2a (turquoise blue frame), expressed in dorsally located Olig2/pMN progenitors (red), limits the dorsal extent of high-threshold Shh response to prevent activation of Nkx2.2a and thus allows maintenance of Olig2 expression (red) required to orient these cells toward the MN lineage. When *sulf2a* is depleted (a’), the range of high-threshold Shh response extends dorsally, leading to dorsal misexpression of Nkx2.2a which in turn represses Olig2. Subsequent changes in the sizes of the p3 and pMN domains cause overproduction of V3 interneurons at the expense of MNs. In 36hpf wild-type embryos (b), the Shh source has expanded in the former Nkx2.2a/p3 domain to form the lateral floor plate (LFP, brown). Immediately adjacent progenitors, i.e. the ventral-most cells of the Olig2/pMN domain, because they activate high-threshold Shh response, upregulate Nkx2.2a. At that stage, Nkx2.2a no more represses Olig2, leading to formation of the p* domain (yellow), populated by progenitors co-expressing Olig2 and Nkx2.2a. Sulf2a expression (turquoise blue frame) is excluded from p* cells but maintained in dorsally located Olig2 progenitors (red) where the enzyme, again, prevents high-threshold Shh response. Then, the two distinct populations of Olig2 progenitors, expressing Nkx2.2a (yellow) or not (red) generate the myelinating (yellow) and non-myelinating (red) OPC, respectively. In 36 hpf *sulf2a*-depleted embryos (b’), LFP formation is not affected but, because of the dorsal expansion of the Nkx2.2a/p3 domain that occurred at earlier stage, the dorsally adjacent progenitors express Nkx2.2a but not Olig2 (green), in contrast to the wild-type situation. Nonetheless, Sulf2a depletion, again, allows activation of high-threshold Shh response that causes dorsal misexpression of Nkx2.2a. Subsequently, the p* domain (yellow), while positioned dorsally, forms properly but, the reduced size of the Olig2/pMN domain leads to a deficit in progenitors that only express Olig2 and, consequently to a strong reduction in the population of non-myelinating OPCs (red). It has to be noted that the population of myelinating OPC (yellow) is also mildly reduced possibly due to a secondary effect of aberrant neuronal populations (see discussion).

The first main finding of our work is that, in contrast to other vertebrates where the different Sulfs display overlapping expression domains in the developing spinal cord ^10,11,24-26^, in zebrafish, a subset of spinal neural progenitors only expresses Sulf2a throughout neurogenesis and gliogenesis. Starting out from this observation, studying the role of Sulf2a in zebrafish represented a unique opportunity to investigate the still debated question of whether Sulf proteins fulfill redundant functions or contribute differentially to spinal cord development^9,11,24^. During the neurogenic phase, which takes place between 10 and 30 hpf in the ventral zebrafish spinal cord^28^, we showed that Sulf2a prevents excess production of V3 interneurons and fosters MN generation. Later on, from 36 hpf, Sulf2a, while not essential for OPC production, has proven to play a key role in triggering generation of non-myelinating OPCs. This markedly contrasts with the function of Sulf1 which stimulates V3 interneuron production, has no impact on MN generation and proves to affect more OPC generation when depleted^7^. Thus, our work makes clear that Sulf2a and Sulf1 exert differential functions in the ventral spinal cord.

We next provide evidence that Sulf2a is part of the regulatory process that provides positional identity to ventral neural progenitors. Specifically, Sulf2a is required, either at neurogenic or gliogenic stage, to maintain a pool of progenitors that express Olig2 but not Nkx2.2a. At neurogenic stages, *sulf2a* expression defines the ventral boundary of the Olig2/pMN domain and acts to prevent dorsal expansion of Nkx2.2a expression within this domain. In agreement with the well-known repressive activity of Nkx2.2a on *olig2* expression at this stage^17,20,31,47^, dorsal misexpression of Nkx2.2a observed in Sulf2a-depleted embryos is accompanied by a marked reduction in the number of Olig2 progenitors. Thus, the progenitor fate change caused by Sulf2a depletion is sufficient to explain the overproduction of V3 interneurons at the expense of MN development observed in these embryos. However, also likely contributing to V3 interneuron overproduction, is the presence of Nkx2.2a/p3 progenitors that persist in Sulf2a-depleted embryos at gliogenic stage. Indeed, in contrast to the wild-type situation, in these embryos, we observed that the Nkx2.2a/p3 domain extends dorsally to the Shh-expressing domain (lateral floor plate).

A somewhat similar but more complex situation was found regarding progenitor identity regulation by Sulf2a at the onset of gliogenesis where *sulf2a* expression, still excluded from the Nkx2.2a-expressing domain, thus from the p* domain, is detected in a subset of Olig2 progenitors. Consistently, the enzyme appears dispensable for formation of the p* domain but required for proper development of progenitors that only express Olig2. How Sulf2a might control development of this particular progenitor population however remains elusive. Since Nkx2.2a no more represses Olig2 at initiation of gliogenesis, dorsal misexpression of Nkx2.2a cannot provide a simple explanation to the loss of Olig2 progenitors caused by Sulf2a depletion as it is the case at neurogenic stage. An alternative explanation might be that Sulf2a plays a role in progenitor recruitment, a process that occurs at initiation of gliogenesis and during which progenitors positioned dorsally to the pMN domain descend to this domain and only then initiate Olig2 expression, a process known to depend on Shh signalling ^41,48^.

Quite apart from the question of how Sulf2a regulates neural patterning, our data shed also new light on the still open question of the origin and relationship of progenitors that give rise to the two OPC subtypes in zebrafish. Two distinct types of spinal OPCs, ones expressing Nkx2.2a and differentiating rapidly as oligodendrocytes and others, not expressing Nkx2.2a and remaining as non-myelinating OPCs have recently being shown to arise from distinct Olig2 progenitor pools ^40,41^. However, how these OPC populations, heterogeneous with respect to Nkx2.2a expression and fate, are generated remained elusive. Our data, showing that Sulf2a depletion preferentially affects generation of both non-myelinating OPCs and Olig2 progenitors that do not express Nkx2.2a, strongly support the view of a lineage relationship between these cells. In apparent contradiction with this proposal, Nkx2.2a-expressing OPCs are also affected, although more moderately, by Sulf2a depletion despite the presence of a correct number of Olig2/Nkx2.2a-co-expressing progenitors. Since neuronal signals and activity can alter OPC development (see ^49^ for review), this result can be potentially explained by the aberrant neuronal populations generated upon Sulf2a depletion.

Finally, examination of Shh signalling activity upon Sulf2a depletion together with the well-known dependence of Olig2 and Nkx2.2a expression on differential Shh thresholds^37,42,43^ are consistent with a role of Sulf2a in attenuating Shh activity at the surface of Olig2 progenitors. Consistently, development of the non-myelinating OPC subset, which is particularly affected by Sulf2a depletion, depends on lower doses of Shh as compared to myelinating OPCs^41^. Our work thus brings support to an essential role of Sulf2a-dependant HS sulfation in regulating Shh signal reception. This is in line with previous reports showing that the sulfation level of glypicans, a family of GPI-anchored HSPGs acting either as co-receptors or as competitors of Shh binding to its receptor Patched, is an important level of regulation of their function at the surface of Shh receiving cells^50-53^. Importantly, this work reveals that Sulf2a plays an opposite role than Sulf1 which stimulates Shh signalling in the developing spinal cord^7-10^. The differential expression of Sulf2a in neural progenitors and of Sulf1 in Shh-producing cells indicates that Sulfs can fulfill opposite functions on the very same signalling cascade by acting either at the source or in receiving cells. This is reminiscent to the dual function of DSulf1, the unique Sulfatase gene in Drosophila, which is expressed both in Hh-producing and -receiving compartments where it exerts a positive and a negative regulatory function on Hh signalling, respectively^54^.

Overall, our work identifies Sulf2a as a crucial regulator of neural cell diversification within the ventral developing spinal cord through a negative action on Shh signalling. Sulf2a contributes to pattern gene expression in neural progenitor cells, sustaining low-threshold response to Shh morphogen at the two critical periods of neuronal and glial production. This work not only unravels a novel function for Sulf2a but also highlights how extracellular sulfatases contribute in different ways to a given signalling pathway depending on their spatial expression in a complex tissue.

## Figure legends

**Supplementary Figure 1: *Sulf2a* depletion impairs motor neuron production**

a-c: Detection (a-c) and quantification (d) of MNs by immunodetection of Islet1/2 (green) in Tg(*olig2*:DsRed) embryos (red) injected with ctrlMO (n=22), *sulf2a*MO^ATG^ (n=13) or *sulf2a*MO^splice^ (n=18) from two independent experiments. Datasets were compared with Mann Whitney’s test (two-tailed). Data are presented as mean number of cells per embryo± s.d (**p< 0.01, **** p<0.0001).

## Methods

### Animals and fish lines

Adult fishes were handled in a facility certified by the French Ministry of Agriculture (approval number A3155501). The project has received an agreement number APAFIS# 2017061313143443 #10204. All efforts were made to minimize the number of animals used and their suffering, in accordance with the guidelines from the European directive on the protection of animals used for scientific purposes (2010/63/UE) and the guiding principles from the French Decret 2013–118.

The following transgenic lines were used to visualise OPCs: *Tg(olig2:*EGFP*)vu12*^*35*^, *Tg(olig2*:dsRed2*)vu19*^*36*^ and *Tg(nkx2.2a:*mEGFP*)vu17*^55^. Shh signalling activity was assessed using the Tg(*GBS-ptch2*:EGFP) transgenic line^44^.

### Generation of *sulf2a* mutant by Crispr/Cas9 technology and genotyping

sgRNAs targeting the exon 11 of *sulf2a* gene was designed using the CRISPOR selection tool ^56^. The following primers were purchased from Integrated DNA Technologies (IDT): 5’ TAGGCTTCCTCTTGAGAGGCGCTG 3’ and 5’ AAACCAGCGCCTCTCAAGAGGAAG 3’, annealed and cloned into a pDR274 host vector (Addgene). After sequence-based selection, this vector was linearized and used as a template for producing RNAs by *in vitro* transcription (T7 mMessage mMachine, Ambion). The obtained sgRNA solution was mixed with Cas9 nuclease (NEB Cas9 Nuclease, S.pyogenes 20mM, M0386M) at 45ng/µl final concentration and used for microinjection in Tg(*Olig2*:GFP) one-cell stage embryos. Founders were identified by sequencing PCR products obtained with the following oligonucleotides: 5’ TTCCTCTGTGCAGAAGTGGC 3’ and 5’ TCAATCTGACAACCTGCGCT 3’. We selected a founder carrying an indel mutation (16 nucleotide deletion and 9 nucleotide insertion) generating a STOP codon at position 529. This F0 founder was outcrossed to generate F1 and phenotypic analyses were performed on subsequent generations by incrossing heterozygous carriers. Genotyping was performed from genomic DNA extracted from tail clipping for adults or from dissected heads from embryos, using the DirectPCR Lysis Reagent (Viagen), supplemented with Proteinase K. PCR fragments were amplified (Go Taq, Promega) using the oligonucleotides mentioned above and subsequently sequenced by Eurofins genomics. Genotypes were interpreted from obtained chromatographs.

### Morpholino injection

Microinjections were carried out in one or two-cell stage embryos using a pressure microinjector (Eppendorf FemtoJet). The following Morpholinos (MOs) (Gene Tools, LLC) were used: *sulf2a*MO^ATG^ (5’-GACCTTAACCAGTGCTCCACCTCCC-3’, 80µM), *sulf2a*MO^splice^ (5’-GAAACTATACTGTGACCCACTCTTC-3’, 1000µM) and standard control MO targeting the human β-globin (ctrlMO, 1000µM). Embryos were injected with a 0.1nl solution. Data presented for each MO are combined for at least two independent experiments.

### Staining procedures

#### Fixation and storage

Embryos were raised and staged according to standard protocols ^57^. Staged embryos were released from the chorion using forceps and placed in a 3.7% formaldehyde solution overnight at 4°C for *in situ* hybridisation or 1h at room temperature for immunostainings. Then embryos were dehydrated through a methanol series and stored in 100% methanol at –20°C.

#### Immunodetection

After rehydration, embryos were treated with 0.25% Trypsin/EDTA (Sigma) at 37°C from 1 to 7 minutes according to their developmental stage. Subsequent incubations and washes were performed in PBS containing 0.8% Triton. The following primary antibodies were used for an overnight incubation at 4°C: rabbit anti-Sox10 (1/1000, Genetex), rabbit anti-Sox2 (1/1000, Genetex), mouse anti-Islet1-2 (1/10, DHSB, 39.4D5). GFP and DsRed2 expression in transgenic embryos was detected using chick anti-GFP (1/1000, Abcam) and rat anti-RFP (1/1000, Chromotek) primary antibodies, respectively. Subsequently, Alexa fluor 488, 555 and 647-conjugated secondary goat antibodies (anti-chicken antibody was purchased from Abcam and all others from Molecular probes) were used at 1/500 for 6 hours at room temperature. For triple immunostainings, we sequentially performed double and single immunodetections. Embryos were mounted in mowiol for subsequent observation.

#### Wholemount in situ hybridisation

The following RNA probes were used: *sulf2a* and *sulf2b* ^*27*^, *olig2* ^37^, *shh* ^58^, *sim1a* ^59^. In all cases, washes were performed in PBS containing 0.5% Tween (PBT) and the hybridisation step was achieved over 24h.

##### Single *in situ* hybridization

was adapted from Xu et al. ^60^ with substantial changes in the hybridisation solution. Briefly, rehydrated embryos were permeabilized with a proteinase K treatment (10μg/ml) from 1 to 45 minutes according to developmental stage, washed in PBT and then re-fixed 20 minutes in a 3.7% formaldehyde/0.1% glutaraldehyde solution. After PBT washes, embryos were placed 3 hours at 65°C in the hybridisation mix containing 50% formamide, 5% Dextran, 1.,3X SSC pH5. 5mM EDTA pH8, 50µg/ml tRNA yeast, 0.2% Tween, 0.5% CHAPS, 0.1mg/ml heparin. Then, hybridisation mix containing the diluted digoxygenin-labelled probe was applied over 24 hours at 65°C. Subsequently, embryos were washed several times in hybridisation mix at 65°C. After a wash in a 50% dilution of hybridisation solution in TBST (25mM Tris Base, 140mM NaCl, 2.7mM KCl, 0,1% Tween) at 65°C, immunodetection was achieved with anti-digoxygenin antibody (Roche) diluted at 1/2000 in TBST/2% Blocking reagent (Roche)/20% Goat serum. NBT/BCIP color reaction was started after TBST washes and monitored under a dissecting microscope. Embryos were mounted in mowiol for subsequent observation.

##### Double fluorescent *in situ* hybridization

Protocol was adapted from Denkers et al. ^61^. After proteinase K permeabilisation, embryos were post-fixed 30 minutes with a 3.7% Formaldehyde solution at room temperature. Fluorescein and digoxygenin labelled probes were incubated together and revealed sequentially: HRP-conjugated antibody against fluorescein (Roche, 1/2000) was incubated overnight and revealed with Cyanine3 TSA (Perkin Elmer). Subsequently, peroxidase was inactivated by 45 minute incubation in 1% H2O2. After 3 washes, HRP-conjugated antibody against digoxygenin (1/1000; Roche) was applied overnight and revealed with 488-TSA solution (Perkin Elmer). Embryos were mounted in mowiol after at least 1 hour of PBT washes.

##### Fluorescent *in situ* hybridisation coupled with immunodetection

was adapted from Denkers et al. ^61^. After hybridisation with the *olig2* digoxygenin-labelled probe, HRP-coupled anti-digoxygenin antibody (1/1000, Roche) was incubated together with chick anti-GFP and rabbit anti-Sox2 primary antibodies overnight. Subsequently, Alexa 488-conjugated goat anti-chick and Alexa 647-coupled goat anti-rabbit secondary antibodies were applied 5 hours at room temperature. After PBT washes, detection of the HRP-coupled antibody was achieved by applying Cyanine3 TSA reagent (Perkin Elmer) 30 minutes at room temperature. Embryos were mounted in mowiol after at least 1 hour of PBT washes.

### Sectioning

After staining, embryos were prepared as described in Andrieu et al. ^62^. Briefly, embryos are incubated overnight in Phosphate Buffer (PB)/15% sucrose at 4 °C. Then, embryos were incubated 2 hours in PB/15% sucrose/7.5% gelatine at 42 °C and transferred into a in PB/15% sucrose/7.5% coated dish. Individual blocks were prepared under a dissecting microscope, frozen in isopentane (Sigma, 615838) at –80°C and stored at –80°C. Cryostat Leica CM1950 was used to generate 20μm thick sections. Slices were then incubated 20 minutes in PB at 42°C to remove gelatin and mounted in mowiol for subsequent observation.

### Imaging, cell counting and statistical analysis

*In situ* hybridisation images were collected with Nikon digital camera DXM1200C/Nikon eclipse 80*i* microscope. Fluorescence photomicrographs were collected on Leica SP8 confocal microscope and were always represented as single optical plane sections. Images were processed (size adjustment, luminosity and contrast, and merging or separating layers) using Adobe Photoshop CS2 (Adobe). In all experiments, cell counts were performed in a defined window positioned along the antero-posterior axis of the spinal cord between somites 14 and 18. Provided data are the average of at least nine embryos per condition from at least two independent experiments. Numbers of individuals (n) are indicated in figure legends. Cell counts of *sim1a*+ cells detected by *in situ* hybridisation in Figure 2 were performed by direct observation under the microscope using a 40x objective, within a 300µm window. For fluorescent stainings, cell counts were performed from z-stacks collected on the entire spinal cord depth and using the Multipoint tool on Image J software. Cell counts of OPCs in Figures 3 and 6 were performed in a 212 µm window. Motor neurons in Supplementary Figure 1 were counted in a 123 µm window. To get information on the dorso-ventral extension of progenitor domains (Figures 4, 5 and 8), we adapted a specific cell counting method. On a z-stack encompassing a 145 µm window, straight lines, distant from 20 µm, were drawn perpendicular to the antero-posterior axis of the spinal cord. Along each of these lines, neural progenitors, marked by Sox2 expression, were counted. Counts were performed on only one side of the lumen which is easily recognisable on z-stacks. For each embryo, the 5 cell counts were used to calculate an average value which was further used to perform statistical analyses. Statistical analyses were performed using Prism5 software (GraphPad). Normality of data sets was tested using Kolmogorov-Smirnov’s, D’Agostino and Pearson’s and Shapiro-Wilk’s tests using Prism5 (GraphPad). We considered datasets as normal when found normal in the three tests. Datasets following a normal distribution were compared with Student’s *t*-test (unpaired two-tailed) while datasets that did not follow a normal distribution were compared using Mann Whitney’s test (two-tailed). For each experiment, the statistical test used is indicated in figure legends. Statistical significance was accepted at a p < 0.05. All data are expressed as mean number of cells per embryo± standard deviation (s.d).

## Acknowledgements

We are grateful to Developmental Studies Hybridoma Bank (Iowa City, IA, USA) for supplying monoclonal antibodies. We thank B. Appel, S. Temple, U. Strahle, C. Winkler, P. Blader, C. Houart and R. Karlstrom for sharing transgenic lines and reagents, E. Theveneau and B. Benazeraf for critical reading of the manuscript; A. Laire and the animal facility staff for zebrafish care and the Toulouse Regional Imaging Platform for technical assistance in confocal microscopy. Work in C.S.’ lab was supported by grants from Agence Nationale de la Recherche (ANR-15-CE16-0014-02), Centre National de la Recherche Scientifique (CNRS), Fondation pour l’Aide à la Recherche sur la Sclérose en Plaques (ARSEP), Fondation ARC pour la Recherche sur le Cancer and Université de Toulouse.

## Authors’ contributions

CD planned the experiments. CD, RD, NE, VB, DO and BG performed and analysed experiments. PC participated in imaging the data. CD and CS supervised the project and wrote the manuscript. All authors read and approved the final manuscript.

## Additional information

The authors declare that they have no competing interests.

