## Supplementary figures and images for "Sulf2a controls Shh-dependent neural fate specification in the developing spinal cord"

### Supplementary Figure 1

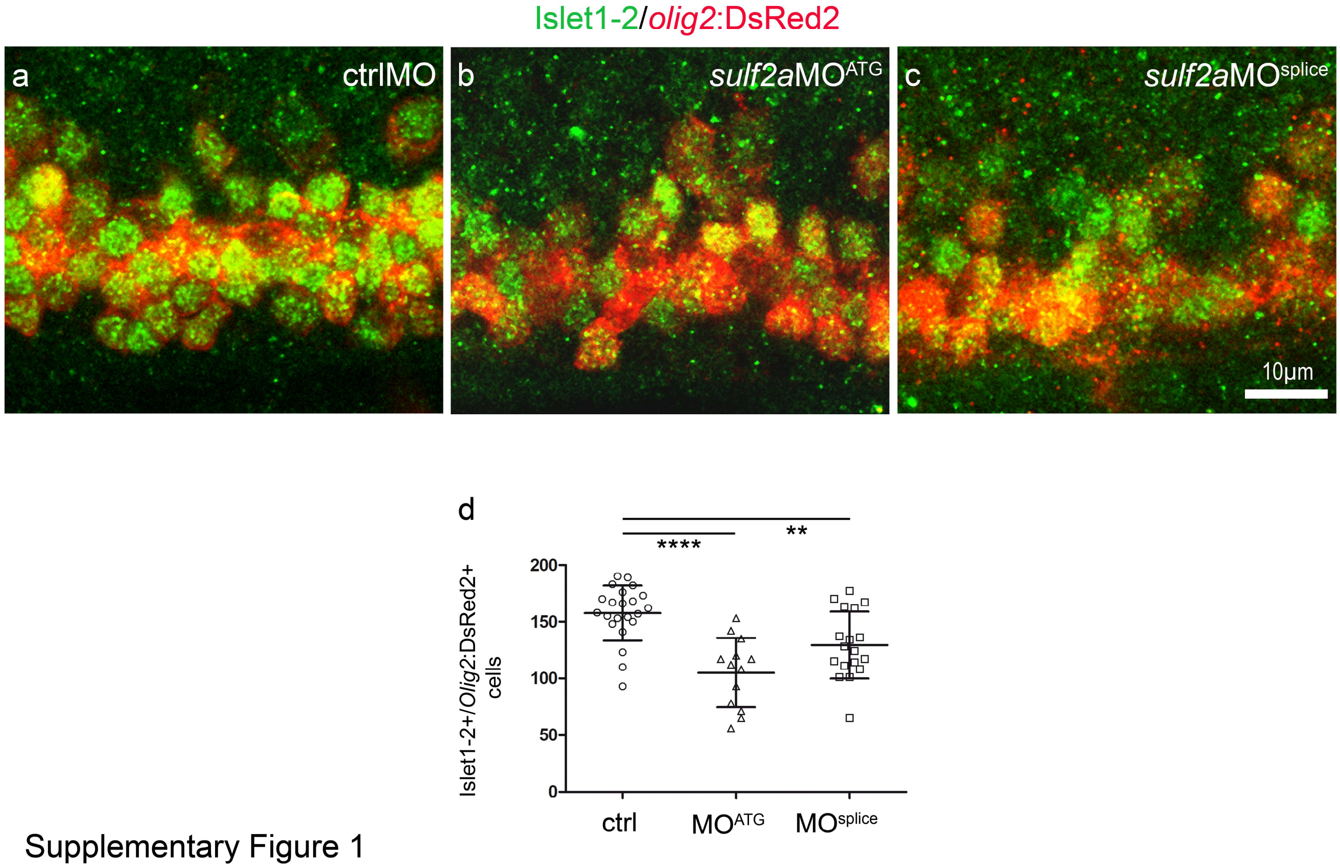


Supplementary Figure 1
